# An Automated Open-Source Workflow for Standards-Compliant Integration of Small Animal Magnetic Resonance Imaging Data

**DOI:** 10.1101/667980

**Authors:** Horea-Ioan Ioanas, Markus Marks, Clément M. Garin, Marc Dhenain, Mehmet Fatih Yanik, Markus Rudin

**Affiliations:** Institute for Biomedical Engineering, ETH and University of Zurich; Institute of Neuroinformatics, ETH and University of Zurich; Centre National de la Recherche Scientifique (CNRS), Université Paris-Sud

**Keywords:** fMRI, MRI, standardization, repositing, data structure, small animal imaging, mouse, rat, mouse lemur, rodent, primate, biomedical engineering, FOSS, BIDS, Bruker, ParaVision, operator guidelines, Python, Bash

## Abstract

Large-scale research integration is contingent on seamless access to data in standardized formats. Standards enable researchers to understand external experiment structures, pool results, and apply homogeneous preprocessing and analysis workflows. Particularly, they facilitate these features without the need for numerous potentially confounding compatibility add-ons. In small animal magnetic resonance imaging, an overwhelming proportion of data is acquired via the ParaVision software of the Bruker Corporation. The original data structure is predominantly transparent, but fundamentally incompatible with modern pipelines. Additionally, it sources metadata from free-field operator input, which diverges strongly between laboratories and researchers. In this article we present an open-source workflow which automatically converts and reposits data from the ParaVision structure into the widely supported and openly documented Brain Imaging Data Structure (BIDS). Complementing this workflow we also present operator guidelines for appropriate ParaVision data input, and a programmatic walk-through detailing how preexisting scans with uninterpretable metadata records can easily be made compliant after the acquisition.

## Introduction

Magnetic resonance imaging (MRI), and functional MRI (fMRI) are highly popular methods in the field of neuroscience. Their high tissue penetration makes them eminently suited for reporting features at the whole-brain level *in vivo*. High assay coverage is particularly relevant for an organ as holistic in its function as the brain, as it facilitates the interrogation of not only sensitivity but also regional specificity. However, MRI methods generate signal via nuclear spin polarisation — which is commonly very weak — and characteristically posess low intrinsic sensitivity. Additionally, fMRI methods rely on highly indirect measures of neuronal activity, and are consequently susceptible to numerous confounding factors.

In animal fMRI in particular, subject preparation, and more specifically cerebrovascular parameters [1] and anesthesia [2, 3] are widely known drivers of result variability. In order to integrate data which may be thus strongly confounded — as well as in order to clarify the confounds themselves [4] — it is vital that data is shared in a raw state, i.e. having undergone no or as little processing as possible. Raw data sharing increases transparency and reproducibility, as data can be assumed to be free from undocumented “fixes”. Such attempts at *ex post facto* data improvement may not just include data matrix manipulations, but also outlier (subject or session) filtering. While valid rationales for both outlier filtering and data editing exist, these processes are best performed in a transparent and well-documented fashion, leaving the raw data untouched as an ultimate recourse.

As published data is intended for reuse, it is reasonable to assume that it may be employed to explore hypotheses other than those under the constraints of which it was originally acquired. In such cases, it is vital that the shared entry and feature pool be as inclusive as possible. It is likely, above all in the effort of methodological comparison and improvement, that what is an artefact or outlier for the interrogation of a narrow hypothesis, may constitute a strong driver of the effect of another hypothesis. Therefore, it is the best choice for small animal MRI researchers to publish data in as raw a form as possible.

The documentation of directed information processing is known as data provenance, and in the effort of establishing a point of recourse it is helpful to map out data traces to the earliest record or earliest record in digital form. In fig. 1, a simplified summary of data provenance is showcased, based on the most common features of small animal MRI. While the rawest data is in theoretical terms the best possible recourse, the extent of this overview illustrates that the choice of a raw data origin point is also constrained by the scope of a researcher’s work. Particularly the first step, reconstructing volumetric information from the time domain k-space record, is commonly not covered by modern MRI pipelines, and instead left to the original acquisition software.

**Figure 1:**
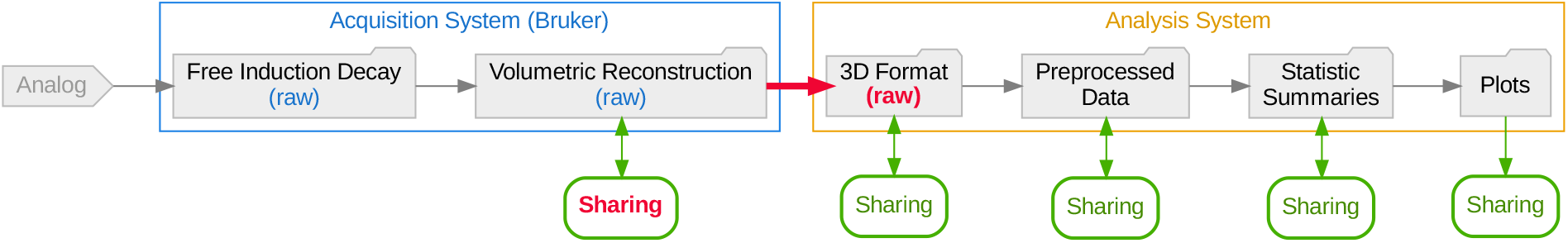
Data provenance flowchart, with the leftmost data being the rawest. Folder nodes represent data states on disk, with nodes suitable as raw data recourse highlighted with the word raw in parentheses. The introduction of a standardized conversion process (red arrow edge) would permit the creation of a 3D Format representation usable as raw data recourse, as well as the sharing of Bruker acquisition system volumetric reconstruction data (also highlighted in red).

Whether directly from the k-space file or from the reconstructed image, the data needs to be converted into a standard, vendor-independent form. Standards are a cornerstone of scientific collaboration, as they concomitantly enable result comparability and data integration. They are, however, at once beneficial and potentially restrictive, since a standard may impose artificial limitations on usability. Such limitations may materialize in negative restrictions on software tools (e.g. proprietary standards may preclude data processing with non-proprietary tools), restrictions on hypotheses (e.g. data organized by one hierarchical principle may lead to loss or obfuscation of certain category correspondence relationships), or positive restrictions of technologies (e.g. data available in highly specific formats might require usage of and familiarity with highly specialized tools) — with the latter restriction being particularly relevant in database standards.

In order to access the benefits and mitigate the pitfalls of standards compliance, data should be migrated to a form which is openly and thoroughly documented, and easily accessible for user manipulation. In the field of small animal MRI, the vast majority acquisition devices (80 %) are produced by the Bruker Corporation, and thus the largest segment of data is initially formatted according to the ParaVision standard. This standard is largely transparent, with most metadata stored in plain-text files. The data, however, are stored in a binary format, which strongly diverges from the *de facto* standard of NIfTI [5], and for which extensive documentation is not openly available. Conversion tools from the ParaVision standard to NIfTI exist [6, 7], and have more recently also been made available from the manufacturer (presently only in closed-source form and only in ParaVision 360, without backward compatibility). Contingent on the scope limitations of the NIfTI format itself, these tools can however not repackage the majority of metadata represented in ParaVision standard plain-text files. This situation exemplifies how the utility of standards is not only contingent on their suitability for use as a common origin format, but also on their flexibility to accommodate all relevant information.

The Brain Imaging Data Structure (BIDS) standard [8], is a prominent candidate for repositing small animal magnetic resonance imaging data. It is extensively tested and well adopted in the field of human MRI, and its extensible and permissive nature makes it easily adaptable to small animal data — as well as generally accommodating for broad swathes of eclectic use cases. The standard builds upon the NIfTI standard, and is organized around a simple directory hierarchy, with key metadata fields captured in the file and path names. This makes BIDS data intuitive to access from both a console and a graphical user interface. Additionally, the standard’s usage of plain-text metadata files makes it accessible to ubiquitous, minimal, and Free and Open Source (FOSS) tools, e.g. the Standard GNU Utilities, or equivalent core utility implementations.

Given varying specifications, it is common for standards to not map fully onto each other (e.g. one metadata field may not have a clear correspondence relationship in both standards). As data conversion is always based only on the most recent parent format, this means that the risk of data (or more specifically, metadata) loss or obfuscation grows with each transition to a new standard. Thus, in the example of fig. 1, collaborative potential is best served if both the “3D Format” representation is infused with sufficient and sufficiently accessible metadata, *and* the original “Volumetric Reconstruction” is rendered shareable.

Both these goals can be attained by the introduction of an automated open-source workflow which can perform the standard transition. As such, all metadata fields which are identified as equivalent between the ParaVision and BIDS standards can be made accessible in the final form. Conversely, if ParaVision standard data is automatically interpretable as input for a concatenation of processing workflows, this original form can also serve as a shareable raw data recourse.

## The Workflow

The workflow, entitled `bru2bids` (Bruker ParaVision to BIDS), is distributed as part of SAMRI [9], a free and open source workflow package of the ETH and University of Zurich Institute for Biomedical Engineering. This repositing workflow can be used stand-alone, but also serves as a gateway to all the further workflows included in the SAMRI package (encompassing dedicated solutions for all analysis steps showcased in fig. 1). As such, the ParaVision-to-BIDS workflow not only permits users to convert data into a format which is more widely supported and flexible, but also easily links to reference implementations for BIDS-based small animal processing functionalities (e.g. registration [10]).

The workflow reposits data and metadata from the ParaVision standard into a BIDS-compliant form. This process includes the conversion of data from ParaVision volumetric reconstruction files (`2dseq`) to NIfTI files. Additionally, metadata is assigned from the specific ParaVision text files to either the NIfTI header, the BIDS metadata files, or the BIDS directory hierarchy, as applicable. A simplified overview of this process is presented in fig. 2a. A more extensive break-down of metadata sourcing — showing the actual input and output files, and highlighting metadata fields represented in the data paths — is laid out in fig. 2b.

**Figure 2:**
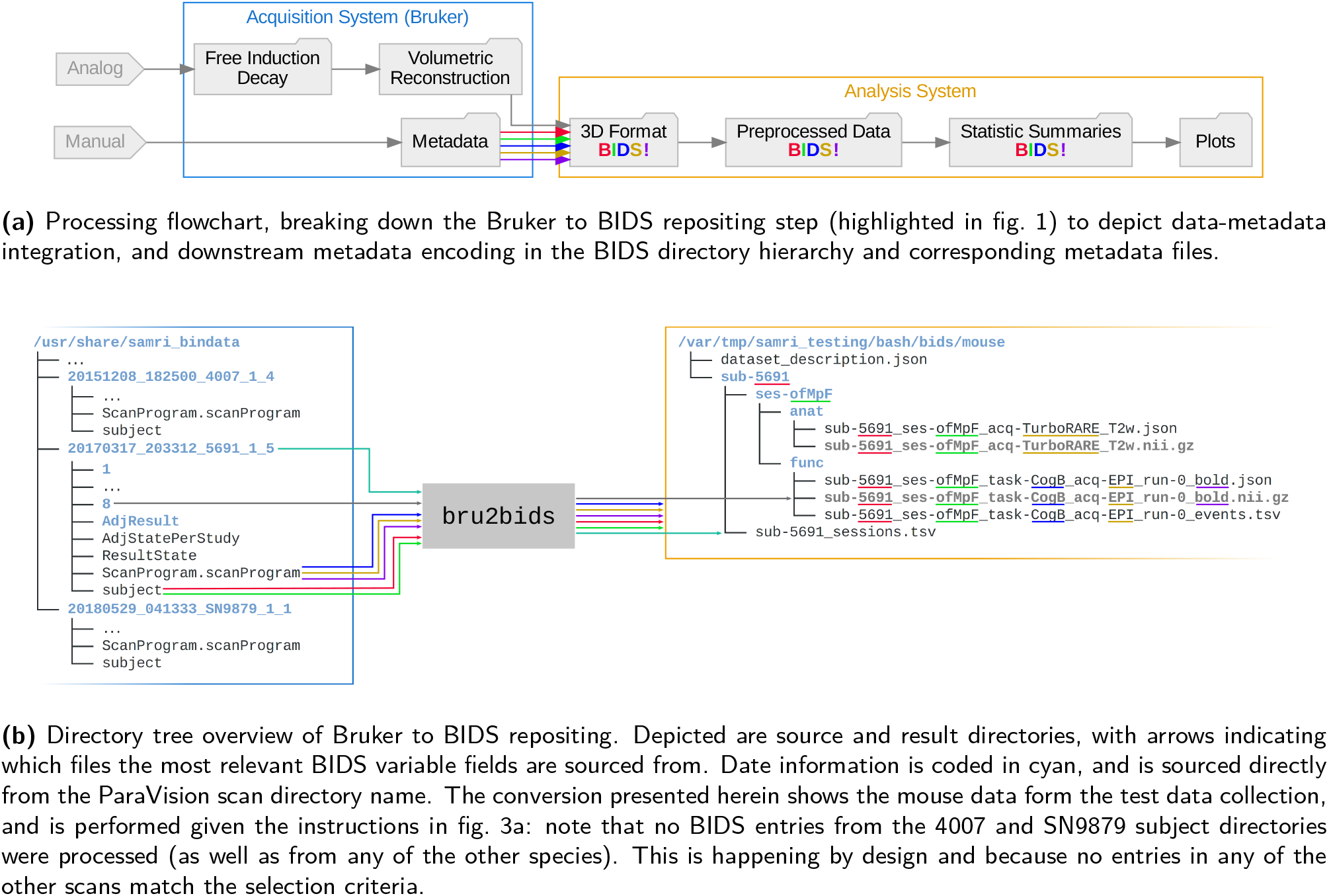
The Paravision-to-BIDS repositing process automatically interprets the source data structure and determines corresponding variables in the BIDS standard. Depicted are color-coded overviews of data and metadata streams during the SAMRI `bru2bids` repositing process, with the data matrix content coded in gray, the subject field in red, the session field in green, the task field in blue, the acquisition field in ochre, and the modality suffix in purple.

The repositing functionality described herein can be accessed from both Bash and Python, via SaMrI bru2bids or `samri.pipelines.reposit.bru2bids()`, respectively. Invocation variants are illustrated in fig. 3, and link to the same code implementation. The `bru2bids` function is highly parameterized, with the same parameter set available in either Bash or Python. A full list of parameters can be obtained by executing the SaMrI bru2bids ---help command from the console. The current parameter listing for Bash is presented under fig. S1.

**Figure 3:**
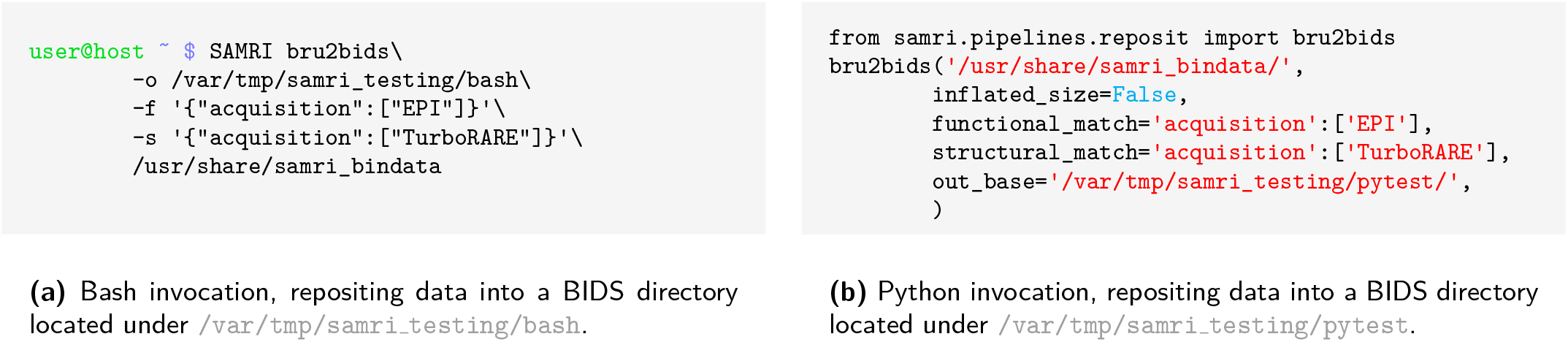
Both Bash and Python can be used to access the repositing functionality. As the Bash binding is auto-generated from the Python function, features become available synchronously, and inspection can be coherently performed regardless of the invocation language. Both code snippets specify the exact same instructions regarding the data source: they indicate that ParaVision standard data from `/usr/share/samri bindata/` is to be reposited into a BIDS standard form, categorizing scans with an “EPI” acquisition string as functional, and files with a “TurboRARE” acquisition string as structural. Both of the above invocations are included in the package test suite.

Notable parameters include the functional, diffusion-weighted, and structural scan specification. These three selectors each use an input dictionary (pairs of one *key*, which is a BIDS metadata string, and a list of accepted *values*), to identify which scans are to be reposited. Examples of such dictionaries are given in fig. 3, where scans with an “EPI” acquisition field are categorized as functional, and files with a “TurboRARE” acquisition field are categorized as structural. The repositing pipeline is run sequentially for all scan categories, as separate processes are required for each scan type (examples of the internal processing nodes are shown in fig. 6).

### Operator Guidelines

The process of scan categorization and metadata sourcing for BIDS conversion is contingent on the presence of operator input records interpretable by the workflow. As the ParaVision metadata files contain free-field input, adherence to a minimal set of guidelines is necessary to ensure unambiguous error-free conversion.

The acquisition of data in the Bruker ParaVision graphical user interface is commenced by creating a new study in the “Study Registration” window. In this window the operator should fill in the “Animal ID” entry corresponding to the intended BIDS subject identifier (i.e. the sub field in the resulting path and file names), and the “Study Name” entry corresponding to the intended BIDS session identifier (i.e. the `ses` field in the resulting path and file names). Notably, the values for both of these fields should be BIDS-compliant — meaning that they should contain only alphanumeric characters, necessarily excluding underscores and hyphens, which are used in the BIDS standard as field separators.

Once the study is created, scans should be renamed in the graphical user interface (before or after acquisition) to contain the additional relevant metadata information according to the BIDS standard. Thus, a resting-state scan acquired with an EPI sequence resolving BOLD contrast should contain the following string in the “Instruction Name” column of the ParaVision interface: `acq-seEPI task-rest bold`. The format is composed out of the BIDS short identifier (e.g. `acq` for acquisition), followed by a hyphen and the desired metadata field value (e.g. `seEPI` for spin-echo echo-planar imaging). The level of detail in these fields is at the discretion of the operator, as per the flexible nature of the standard. If recognizing the EPI variant at-a-glance is deemed irrelevant for the data at hand, this field may simply be assigned a value of `EPI`, or could conversely be expanded to include an arbitrary amount of additional detail. Pairs of BIDS short identifiers and desired values should be separated by underscores, with the modality suffix appended at the end after a final underscore separator.

Additional BIDS fields such as the run ordinal number (e.g. `run-0`, as seen in fig. 2b), are automatically determined by the `bru2bids` workflow. The modality suffix, if not explicitly specified by the operator, can also be automatically assigned in a small number of cases where it is unambiguous (e.g. a scan with a FLASH acquisition but with no operator defined modality will be assigned a `T1w` modality suffix, and similarly, one with a TruboRARE acquisition will be assigned a `T2w` suffix).

Entering metadata in this fashion during scanner operation creates records which can directly serve as input for analysis workflows, and are thus immediately ready for sharing or analysis. Examples for such interpretable metadata scans are available in the SAMRI ParaVision testing data archive, and the example strings specified in this section are sourced from the rat data identified in the archive by the `20171106_184345_21_1_1` directory name.

### Preexisting Data Compliance

Preexisting data sets — acquired with operator metadata input divergent from the recommended guidelines — can also be redressed, via a small number of plain text editing operations. Performing such edits is highly recommended, as it is still easier than manually repositing data in the BIDS standard, and also permits direct ParaVision data sharing, as shown in fig. 1.

The relevant parts of a ParaVision scan directory which need to be edited are single lines in the `subject` and `Scanprogram.scanProgram` files. Editing the subject identifier in the ParaVision directory name is also advisable, though only for ease of overview — since the subject field is not read from the directory name by the repositing workflow.

Before editing, value names for all available metadata fields must be chosen, such as respect the constraints of the BIDS standard. This means that subject, session, task, and all other desired identifiers need to contain only alphanumeric characters (digits 0 through 9, and lower and uppercase letters). Once all identifiers are properly chosen, they can be replaced or retroactively entered into the ParaVision metadata fields.

The `subject` file defines both the subject and the session of the scan. To enter the subject identifier, the line *below* the line containing the string `##$SUBJECT id=` must be edited. This following line should contain the subject identifier between greater than and less than characters, e.g. for a subject identified as Mc365A, the line should read `<Mc365A>`. Analogously, to enter the session identifier, the line *below* the line containing the string `##$SUBJECT study name=` needs to be edited to read e.g. `<R02>` for a session identified as R02 within the study, as shown in fig. 4b.

**Figure 4:**
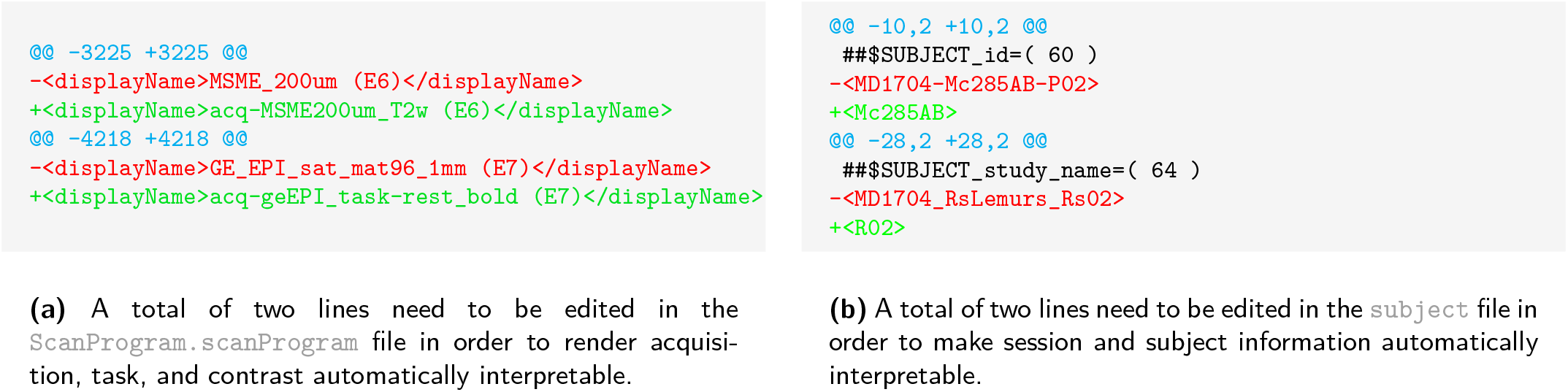
Editing operations on only up to four lines are required to render preexisting data compatible with repositing via the workflow at hand. Depicted are file differences for the `20171024_165248_MD1704_Mc285AB_P02_1_1` testing dataset scan, in a patch syntax. The line numbers identifying the position of the text segment (before and after editing) are highlighted between arobase characters and in cyan. Deleted and added lines are highlighted in red or green, and prefixed with a minus or plus, respectively. Conserved lines are printed in black.

The `Scanprogram.scanprogram` file is used to define other BIDS metadata fields. Within this file, individual lines containing the string u(`E` (with the open box character representing a space) are used both to record the ParaVision “Instruction Name” and to establish correspondence with the respective numbered ParaVision scan directory. On such lines, after the open `<displayName>` tag, and up to the space character before the open parenthesis, an arbitrary sequence of BIDS short identifiers and value pairs, separated by underscores, can be inserted – as shown in fig. 4a.

Examples for manually redressed data are included in the SAMRI ParaVision testing data archive, and are in their end form indistinguishable from data acquired along the lines of the operator recommendations. In fig. 5 we show the changes which were required to render mouse lemur testing data compliant with the repositing workflow.

**Figure 5:**
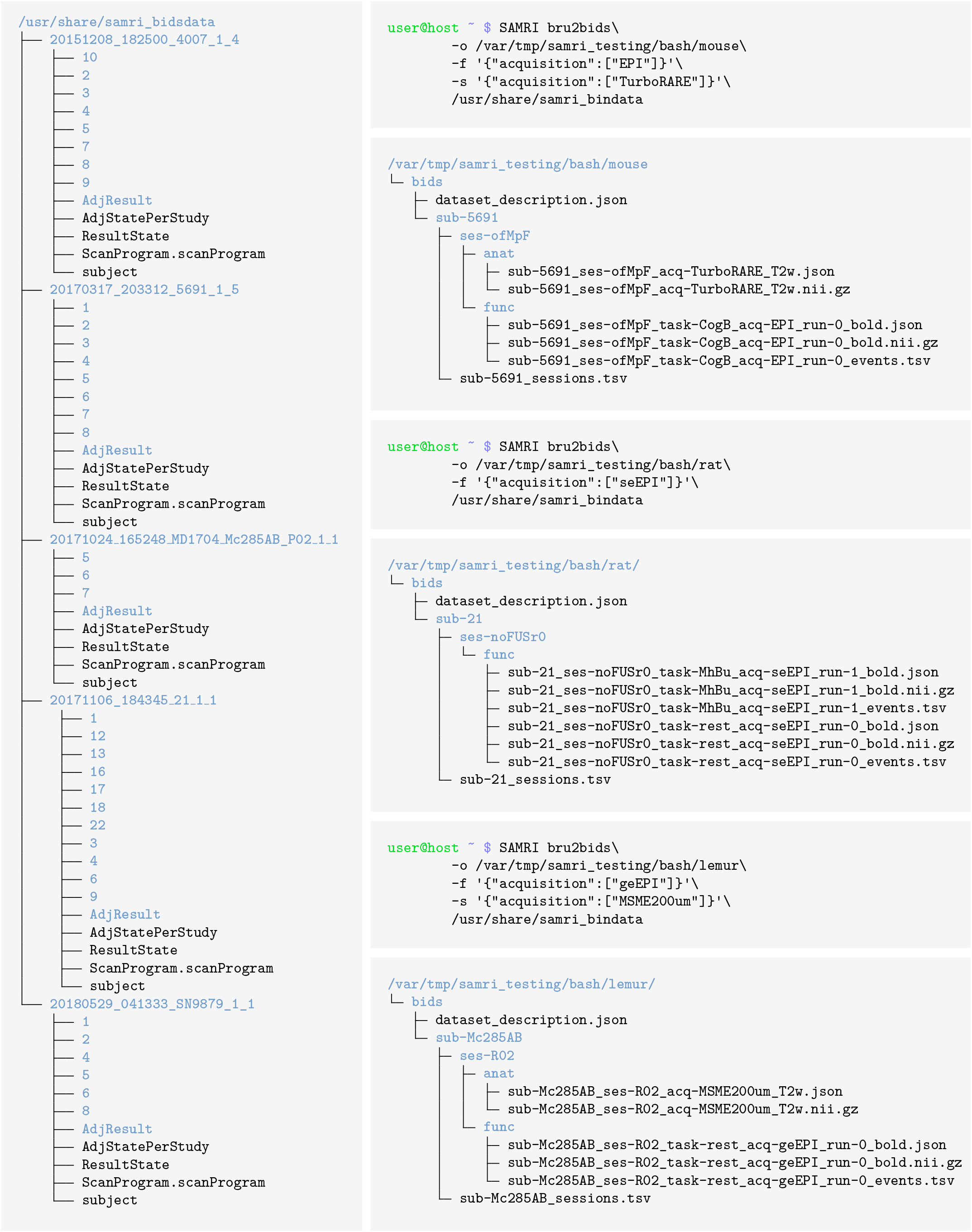
The workflow can automatically select, convert, and reposit ParaVision data from a general collection directory into dedicated shareable data sets, formatted for the BIDS standard. The left column shows the contents of the source directory, analogous to a collection directory on a server or scanner, and packaged by us as `samri bindata`. The right column contains the Bash commands needed to produce dedicated mouse, rat, and lemur data sets (in that order). Below each Bash command shell, the resulting BIDS directory contents are shown. Directories are highlighted in blue. As ParaVision provides no way of integrating stimulatioin information, the workflow creates empty events files (`*_events.tsv`), which the user can fill with the appropriate stimulation content, delete, or simply ignore (if empty the files carry no meaning in BIDS).

## Results

To demonstrate the capabilities of the workflow, we have compiled a versioned reference archive of multispecies small animal MRI data. The resulting package, `samri bindata`[11], includes multi-center ParaVision standard scans from mice, rats, and mouse lemurs, and serves as testing data for the SAMRI package, including the repositing workflow. The data in this archive is based on scans acquired with originally compliant operator input, as well as on preexisting scans rendered compliant *ex post facto*, as detailed in this article.

In fig. 5 we test the capability of the workflow to correctly and automatically source specified datasets from a diverse collection, and reposit them in the BIDS standard. The resulting three separate and shareable BIDS data sets pass the BIDS validator with no errors. The commands listed in this section are included in the SAMRI test suite, and monitored for continued quality assurance.

### Methods for Implementation

The data of our reference ParaVision testing archive, `samri bindata`[11], were acquired at three separate centers, using three different scanner types, and in two rodent species (mouse and rat) and one primate species (mouse lemur) — with ParaVision `6.0.1`. Mouse data were acquired at the Animal Imaging Center of the ETH and University of Zurich, using a Bruker Biospec 70/16 system, or a Bruker Biospec 94/30 system. Rat data were acquired at the Neurotechnology Group of the University of Zurich Neuroinformatics Institute, using a Bruker PharmaScan 70/16 system. Mouse lemur data were acquired at the Molecular Imaging Research Center (MIRCen) of the Commissariat à l’Ènergie Atomique et aux Ènergies Alternatives (CEA), using a 11.7 Tesla Bruker BioSpec system.

The workflow is implemented as a function in the Python programming language, and uses the Nipype package [12] for workflow execution, overview generation, parallelization, and access to non-Python tools. Bash bindings are auto-generated based on the Python function definition and documentation string by the Argh package.

Data conversion from the ParaVision `2dseq` format to NIfTI is performed by the `Bru2` function from the Bru2Nii package [6]. Preliminary to workflow execution, ParaVision metadata parsing and BIDS metadata assignment is performed by Python utility functions implemented in the SAMRI package. These functions iterate through the lines of the relevant metadata files: `subject` and `ScanProgram.scanProgram`, falling back to `<scan number>/acqp` if the scan program file is corrupted. Metadata detection is performed via regular expressions, to afford a maximum of flexibility, and avoid dependency on more convoluted higher-level tools.

The processed metadata is recorded in a Pandas Dataframe [13], which is both used internally and written to disk in the “work directory” to permit debugging. This record is then divided along the lines of the supported scan categories (structural, functional, and diffusion-weighted), and the resulting selections are used to initiate the respective workflow iterations, as seen in fig. 6. Following successful workflow execution, the BIDS standard `*_sessions.tsv` file is separately generated, recording the onset acquisition times for each session.

**Figure 6:**
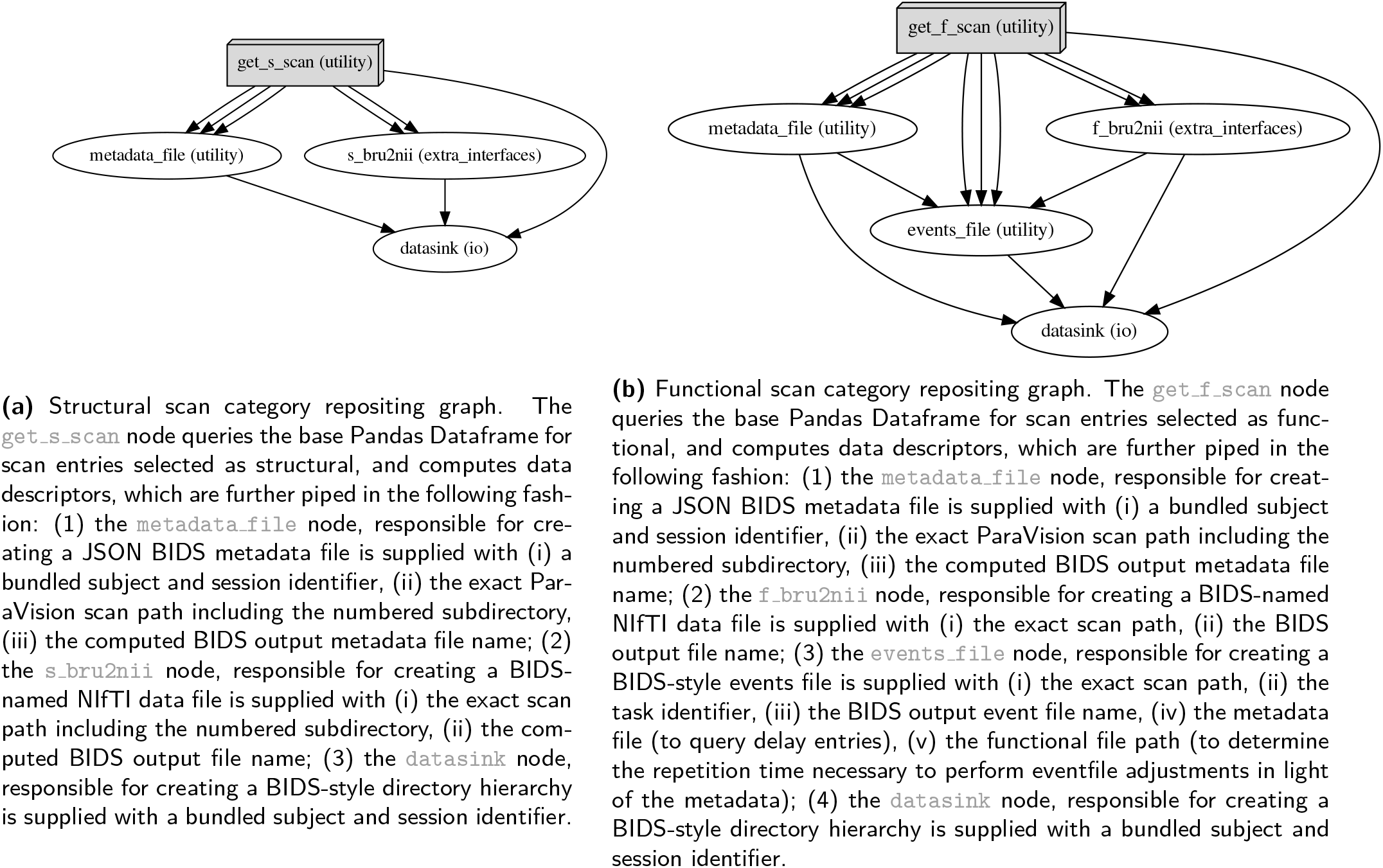
Dedicated workflows are set up for each scan category. Depicted are directed acyclic graphs, as produced by the workflow engine, Nipype. The `bru2bids` function iteratively executes e.g. structural **(a)** and functional **(b)** repositing workflows, contingent on data availability. Node names specify the source code identifiers, with the text in paranteses indicating, which modules they are implemented in: `utility` and `extra_interfaces` are modules of the SAMRI package, and io is a module of the Nipype package.

The guidelines regarding operator input and preexisting data compliance presented in this document are based on the ParaVison `6.x` release series. The parsing functionality of the workflow in the current SAMRI version `(0.3)` is based on the ParaVison `6.x` release series and the BIDS `1.x` specification.

## Discussion

The `bru2bids` workflow presented herein is a significant first step in rendering data in the Bruker ParaVision standard automatically interpretable for high-level analysis pipelines. This is done by repositing the ParaVision data according to the BIDS standard, which offers superior legibility, as well as integration with community analysis tools, specifically with tools adapted from human fMRI. The BIDS reposited data form can serve as a mere intermediary, facilitating data usage with standardized workflows, but can also be used in and of itself as raw data recourse — if data management expediency is prioritized over the larger pool of accessible information in the full ParaVision standard. To demonstrate and persistently track compliance, we release a versioned archive of Bruker ParaVision testing data, diverse in terms of both animal species and acquisition protocols. We test the performance of the workflow on the dataset, and report compliance with the target standard. These demonstrated capabilities of the workflow are rendered accessible to the community by procedural instructions addressed to Bruker MRI scanner operators. Additional accessibility is conferred by a detailed walk-through, which allows custodians of Bruker ParaVision data to render preexisting records compatible as a workflow input. Further, we describe the general principles of the software implementation, which in conjunction with the documentation internal to the software enable collaborators to inspect, debug, augment, or create derivations based on our work.

The demonstrated performance of this workflow, its position at the transition from the most popular small animal MRI acquisition format into the most popular MRI data sharing format, as well as its transparent free and open source nature, make the SAMRI’s `bru2bids` a strong foundation for the rapid and collaborative improvement of fMRI data analysis methodology.

---

**Figure S1:**
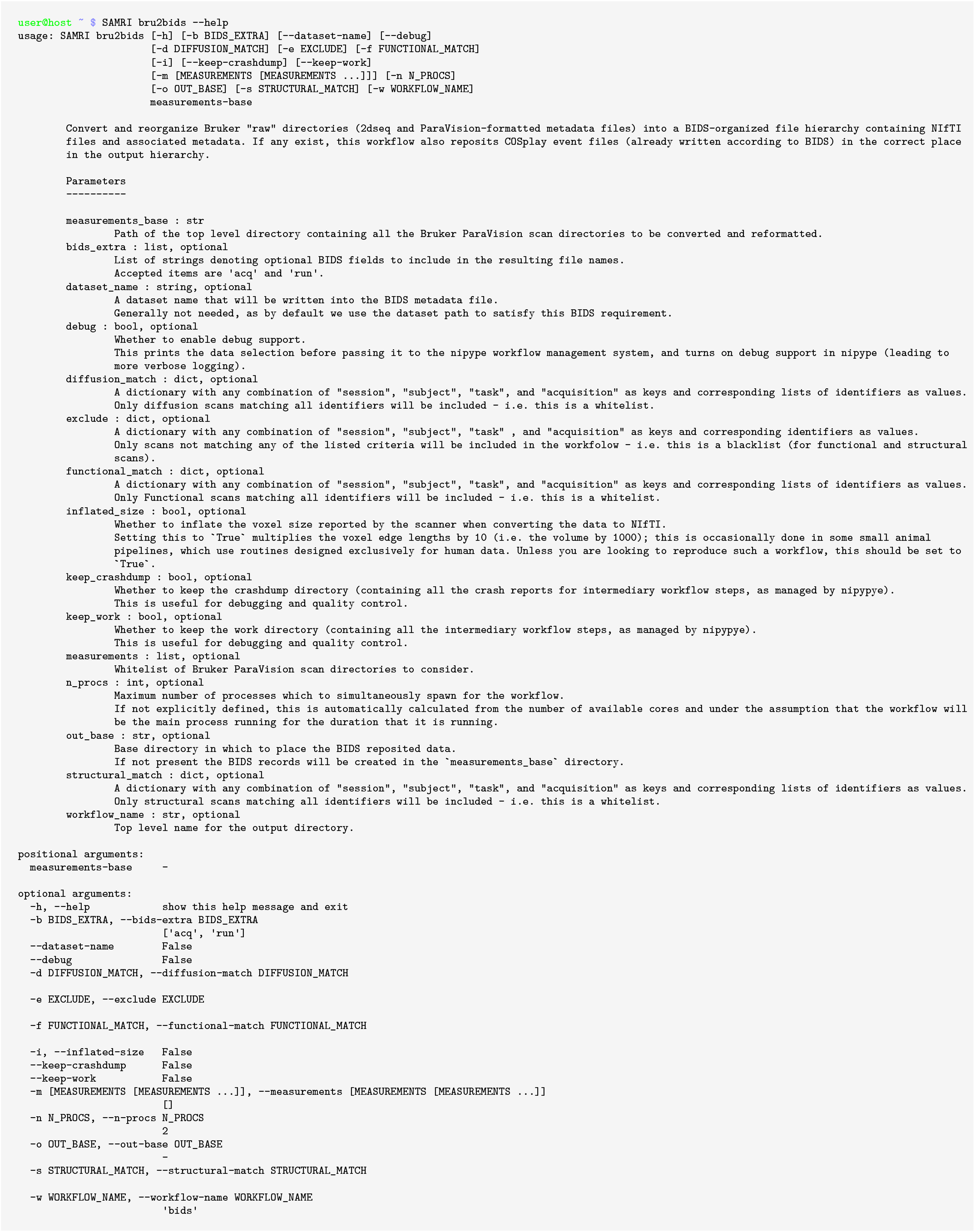
Example of the SAMRI `bru2bids` repositing workflow parameter list as reported in the Bash command line in the `0.3` SAMRI release. This output is autogenerated for Bash from the Python docstring.

**Figure S2:**
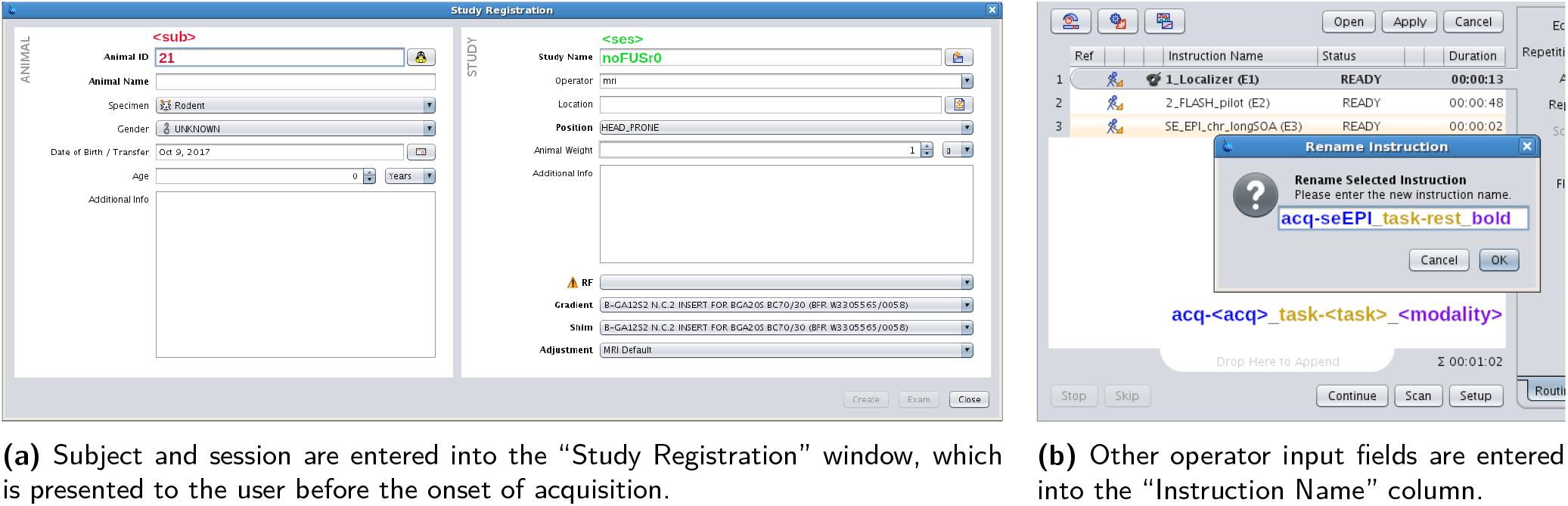
ParaVision graphical user interface examples, showing how metadata automatically interpretable by the repositing workflow can be specified when acquiring new data.

